# Ribosomal protein L23 drives the metastasis of hepatocellular carcinoma via upregulating MMP9

**DOI:** 10.1101/2021.07.27.453993

**Authors:** Minli Yang, Yujiao Zhou, Haijun Deng, Hongzhong Zhou, Shengtao Cheng, Dapeng Zhang, Xin He, Li Mai, Yao Chen, Fan Li, Juan Chen

## Abstract

Hepatocellular carcinoma (HCC) is one of the leading causes of cancer-related deaths globally and tumor metastasis is one of the major causes of high mortality. To identify novel molecules contributing to HCC metastasis is critical to understanding the underlining mechanism of cancer metastasis. Here, combining the analyze based on published database and liver tissues from HCC patients, we identified that RNA binding protein L23 (RPL23) as a tumor metastasis driver in HCC. RPL23 was elevated in HCC and closely related to poor clinical outcomes. Furthermore, RPL23 depletion inhibited HCC cell proliferation, migration and invasion, while RPL23 overexpression promoted HCC cell metastasis. Mechanistically, RPL23 positively regulated MMP9 expression by stabilizing its mRNA. And increased MMP9 is involved in RPL23-mediated HCC metastasis. Importantly, RPL23 silencing reduced tumor growth and metastasis *in vivo*. In summary, we identified that RPL23 play an important role in HCC metastasis in an MMP9-dependent manner and may be a novel potential therapeutic target for HCC tumorigenesis and metastasis.

## Introduction

Hepatocellular carcinoma (HCC) is the predominant malignancy of the liver and ranks as the third most common cause of cancer-related death worldwide ^1, 2^. Despite the innovative progress of HCC management, most are diagnosed at advanced stage when the therapeutic options are limited. As an aggressive malignancy, HCC metastatic spread is the main obstacle to treatment and extension of long-term survival^3, 4^. However, the underlying mechanisms of HCC metastasis have not yet been fully explored. Therefore, deepen the understanding of molecular mechanisms of HCC metastasis is an urgent need to develop novel therapeutic approaches.

RNA-binding proteins (RBPs) are critical regulators of gene expression by which involved in various aspects of RNA metabolism, such as splicing, modification, stability, and translation ^5–8^. Growing number of studies have revealed that altered RNA metabolism due to dysfunction of RBP plays an important role in cancer progression, especially cancer cell metastasis^9, 10^. Notably, dysregulated RBPs were also observed in HCC. Based on the analyses of 1,225 clinical HCC samples, the researchers found that large numbers of RBPs are dysregulated in HCC and is closely related to poor prognosis^11^. SORBS2, a RBP which significantly decreased in HCC, could suppress HCC metastasis by stabilizing RORA mRNA^12^ or inhibiting c-Abl-ERK signaling pathway^13^. Further, elevated RBP Polypyrimidine tract-binding protein 1 (PTBP1) could facilitate HCC tumor growth by interacting with the 5’-UTR of cyclin D3 (CCND3) mRNA to enhance CCND3 translation^14^. While the functions of RBPs in HCC have been extensively investigated, the underlining mechanism of RBP in HCC metastasis is not fully understand. Hence, identification of RBPs in HCC metastasis is necessary to broaden the potential of targeted therapy.

Human ribosomal protein L23 (RPL23), a novel RBP, has been reported that involved in some human cancer progression, such as lung cancer^15^, myelodysplastic syndromes^16^, gastric cancer ^17^, colorectal carcinoma cells^18^. Based on the above report, RPL23 could against tumorigenesis mainly through regulating p53 pathways. As an RNA-binding protein, RPL23 also plays an important role in RNA metabolism. Growing number of studies have revealed that altered RNA metabolism due to dysfunction of RBP plays an important role in cancer progression, especially cancer cell metastasis^19, 20^. However, the effect of dysfunction of RPL23 on HCC metastasis still remains unclear. Thus, the aim of this study is to examine the effects of RPL23 on the metastasis of HCC and to elucidate the underlying mechanisms.

Importantly, matrix degradation plays a critical role in cancer cell metastasis. Matrix metalloproteinases (MMPs) is responsible for the degradation of matrix^21^ which are essential for cell metastasis. MMP9 is one of the most complex MMPs and a group of researchers demonstrated that MMP9 was overexpressed in liver cancer^22^ which involved in HCC metastasis. However, molecular mechanisms of MMP9 upregulation in HCC are largely unknown. It has reported that the expression of MMP9 could be upregulated by RNA binding protein HuR via stabilizing of MMP9 mRNA^23^, indicating that RBPs may involve in MMP9 regulation.

In this study, we first evaluated the expression and prognostic value of RPL23 in HCC, then further explored the effect of RPL23 in HCC migration and invasion by interacting with MMP9 both *in vitro* and *in vivo*. Mechanistically, RPL23 stabilizes MMP9 mRNA and enhances MMP9 expression to promote HCC metastasis. Taken together, we first reported the interaction between RPL23 and MMP9 in regulating HCC metastasis and observed that RPL23 may become a potential effective biomarker for HCC.

## Materials and methods

### HCC tissue samples

HCC tissues and paired adjacent non-tumor tissues were obtained from 60 patients who underwent surgical resection for HCC at the First Affiliated Hospital of Chongqing Medical University in Southwest China, with the approval of the Institutional Review Board of Chongqing Medical University. The patients provided informed consent and had not received any prior radiotherapy or chemotherapy. All specimens were frozen immediately after surgery and stored in liquid nitrogen until use.

### Cell culture

HLE and MHCC97H cell lines were obtained from the Cell Bank of the Chinese Academy of Sciences (Shanghai, China). Huh7 was obtained from the Heath Science Research Resource Bank (HSRRB). Cells were cultured in Dulbecco’s Modified Eagle’s Medium (DMEM), supplemented with 10% fetal bovine serum (Gibco-BRL), 100U/ml penicillin, and 100 μg/ml streptomycin at 37 °C in 5% CO2. All the cells were examined negative for mycoplasma.

### Antibodies, plasmids and chemicals

The complementary DNA of full-length RPL23 was amplified by polymerase chain reaction and inserted into the *pcDNA3.1-Flag* vector at EcoRI and XhoI restriction sites. The MMP9 vector was purchased from OriGene Technologies (Rockville, MD). The specific short hairpin RNA of RPL23 (shRPL23#1:GCAAACCAGCTCAGAAATT, shRPL23#2: GAGTCATAGTGAACAATAATT) were obtained from GenePharm (Shanghai, China). Actinomycin D was purchased from Sigma (SBR00013).

### RT-qPCR analysis

Total RNAs were obtained from the cultured cells or tumor tissues using Trizol reagent (Invitrogen, USA). RNA was quantified by absorbance at 260 nm with a NanoDrop One (Thermoscientific). For RT-qPCR, 1 μg of RNA was reverse transcribed into cDNA using the Reverse Transcription Kit (Bio-Rad, USA). qRT-PCR was carried out using SYBR Green (Roche, Germany), and β-actin was used as control. Primer sequences used for RT-qPCR were listed in Supplementary Table 1.

### Western blot analysis

Cells were washed twice with ice-cold phosphatebuffered saline (PBS), collected in RIPA buffer with protease inhibitor cocktails (Roche, Indianapolis, IN) and lysed on ice for 15 min. Lysates were centrifuged for 5 min at 13 000 × g at 4°C and the concentration of protein was measured using a bicinchoninic acid assay (Thermo Fisher Scientific). Protein lysates were fractionated by SDS-PAGE and transferred onto PVDF membranes. The primary antibodies were as follows: rabbit anti-RPL23 (Proteintech, 16086-1-AP), rabbit anti-MMP9 (OriGene, TA326652), anti-GAPDH (Santa Cruz Biotechnology, sc-365062).

### In vitro assays for migration and invasion

Cell transwell chambers with or without matrigel (24-well plate, 8 m pores; BD Biosciences) were used to assess cell migration and invasion. For migration assay, 1 × 10^5^ HCC cells with different treatments were seeded into the upper chamber of transwells and cultured in serum-free DMEM at 37°C for 18h, while medium with 10% FBS was put into the lower chamber. For invasion assay, indicated cells were seeded in the upper chamber with matrigel-coated membrane. After a certain time, migrated or invaded cells were fixed in 95% methanol and stained with 0.1% crystal violet dye. The number of migrated or invaded cells in five different high-magnification fields (40×) were counted under an inverted microscope.

### Wound-healing assay

For wound-healing assay, equal numbers of HCC cells were plated into six-well plates. Until confluence, the cell monolayer was scratched with a sterile pipette tip to draw a gap, and washed twice with PBS to remove cell debris. Cells were photographed to record the wound width at 0, 24, 48 h, respectively.

### CCK8 assay

For cell proliferation assay, CCK-8 (MedChemExpress, #HY-K0301) assay was used to assess cell viability according to manufacturer’s instructions. Different groups of cells at a density of 3000 cells/well were plated in 96-well plate with 100 μl medium and cultured for 6, 24, 48, 72, 96 h, then maintained in complete DMEM with 10% CCK8 reagent for 2 h at 37 °C. The absorbance at 450 nm was measured by using a plate reader.

### Immunohistochemistry (IHC)

For IHC, HCC tissue samples were fixed in 10% formalin and embedded in paraffin. The tumor tissue sections were prepared, deparaffinized in xylene, rehydrated with alcohol, and then washed in phosphate-buffered saline (PBS). In order to antigen retrieval, tissue sections were heated at 105 °C for 20 min in a citric acid buffer (0.01 M), later dealt with 3% hydrogen peroxide solution to block the endogenous peroxidase activity and blocked with bovine serum albumin (BSA) for 120 min. Next, antibodies of RPL23 and MMP9 were used to incubate at 4 °C overnight, then the tissue sections were incubated with horseradish peroxidase (HRP)-conjugated secondary antibody to detect the target protein. Antigen-antibody chromogenic reactions were developed for 12 min. After that, the slides were stained with hematoxylin and dehydration in graded alcohols and xylene. The immunohistochemical staining was analyzed by Image-Pro Plus 6.0 software.

### Cytoskeletal staining

The different treatment cells were seeded on the glass coverslips and incubated for 24h at 37°C. Media was removed, cells were gently washed for 3 times with PBS and fixed with 4% paraformaldehyde for 10 min at room temperature (RT). Then, cells were permeabilized with 0.5% Triton X-100 for 15 min. After washing with PBS, cells were counterstained with DyLight™ 488 Phalloidin (CST, #12935; dilution 1:40) for 10 min to stain F-actin. Subsequently, cell nuclei were stained with 4′,6-diamidino-2-phenylindole (DAPI). After air drying for 20 min, the cells were sealed with anti-fluorescent quencher. Finally, images were obtained by confocal laser-scanning microscopy using a Laser Scanning Confocal Microscopy (Leica TCS SP2).

### RNA immunoprecipitation (RIP) assay

RIP assay was performed using the Magna RIP RNA IP kit (17–700) from Millipore according to manufacturer’s protocol. In brief, HCC cells (2 × 10^7^) were lysed with RNA immunoprecipitation lysis buffer (Millipore, USA) and then incubated with 2 μg of rabbit polyclonal anti-RPL23 or non-immunized rabbit IgG at 4 °C overnight. The RNA protein immunocomplexes were pulled down by 30ul protein A/G magnetic beads. After RNA purification, qRT-PCR was used to determine the levels of target genes.

### RNA pull-down assay

RNA oligonucleotides labeled with biotin at the 5’-end were synthesized by Integrated DNA Technologies. The RNA sequences used in this study were listed as following, sense-M9-5’UTR-F: AGACACCTCT GCCCTCACC, sense-M9-5’UTR-F: GGTGAGGGCAGAGGTGTCT; sense-M9-3’UTR-F: TAATACGACTCACTATAGGG GGCTCCCGTCCTGCTTTGGC; sense-M9-3’UTR-R: TAAAGGTTAGAGAATCCAAG; M9-CDS-F: TAATACGACTCACTATAGGGATGAGCCTCTGGCAGCCCCT; M9-CDS-R: CTAGTCCTCAGGGCACTGCA. In all, 50 pmol Biotinylated RNA oligos were conjugated with 50 μl of streptavidin beads (50% slurry; Thermo Fisher) in a total volume of 300 μl of RNA-binding buffer (20mM Tris, 200mM NaCl, 6mM EDTA, 5mM sodium fluoride and 5mM β-glycerophosphate, PH 7.5) at 4 °C on a rotating shaker for 2 hours. After washing three times with RNA-binding buffer, RNA-beads conjugates were incubated with 100 μg of nuclear extracts in 500 μl RNA-binding buffer on a rotating shaker overnight at 4°C. The beads were then washed thoroughly three times with RNA-binding buffer, eluted with 30 μl 1 × SDS loading buffer and subjected to SDS-PAGE and western blot.

### Xenograft model of lung metastasis

For in vivo tumor metastasis assays, male BALB/C nude mice (6-8 weeks old) were used for the study. 2 × 10^6^ MHCC97 cells infected with shRPL23 or empty vector were suspended in 40 μL of a 1:1 (v/v) mixture of a serum-free DMEM/Matrigel solution and then orthotopically implanted into the left hepatic lobe of nude mice. After 6 weeks, mice were sacrificed and primary tumor volume and lung metastasis were scored. All of the procedures for handling of animals abided by the guidelines of Chongqing Medical University Animal Care Committee (reference number: 2019002).

### Statistical analysis

The data were presented as mean ± standard deviation. All statistical analyses were conducted using GraphPad Prism8 (GraphPad) software. Unless otherwise indicated, experiments were analysed with a two-tailed Student’s t-test with a confidence interval of 95% when the number of groups equalled 2, or with a parametric ANOVA test when the number of groups was >2. The nonparametric χ2 test was used to assess the correlation between RPL23 expression and the clinicopathological parameters. P <0.05 was defined as statistically significant (*P<0.05; **P<0.01; ***P<0.001).

## Results

### Upregulated RPL23 expression is correlated to poor clinical outcomes in HCC

Based our previous RNA-sequencing data from 10 pairs of primary HCC tissues with extrahepatic metastasis (EHMH) and 10 pairs of metastasis-free HCC tissues (MFH), ribosomal protein L23 (RPL23) is extremely upregulated in HCC tissues, especially in those tissues with extrahepatic metastasis (EHMH). To broadly investigate the potential function of RPL23 in HCC, the expression of RPL23 was first analyzed in 2 published datasets, The Cancer Genome Atlas Cohort (TCGA) and Gene Expression Omnibus (GEO). As expected, RPL23 expression increased obviously in HCC tissues than that in normal tissues (Fig.1A-B, *p*<0.05). Moreover, RPL23 overexpression was clearly associated with advanced tumor grade (Fig.1C, *p*<0.0001) and late cancer stage (Fig.1D, *p*<0.0001). Kaplan-Meier analyses showed that patients with higher levels of RPL23 had shorter disease-free survival (DFS; Fig.1E, HR=1.4, *p*=0.045) and overall survival (OS; Fig.1F, HR=1.6, *p*=0.0069), suggesting that upregulation of RPL23 was closely related to poor clinical outcomes in HCC.

**Figure 1.**
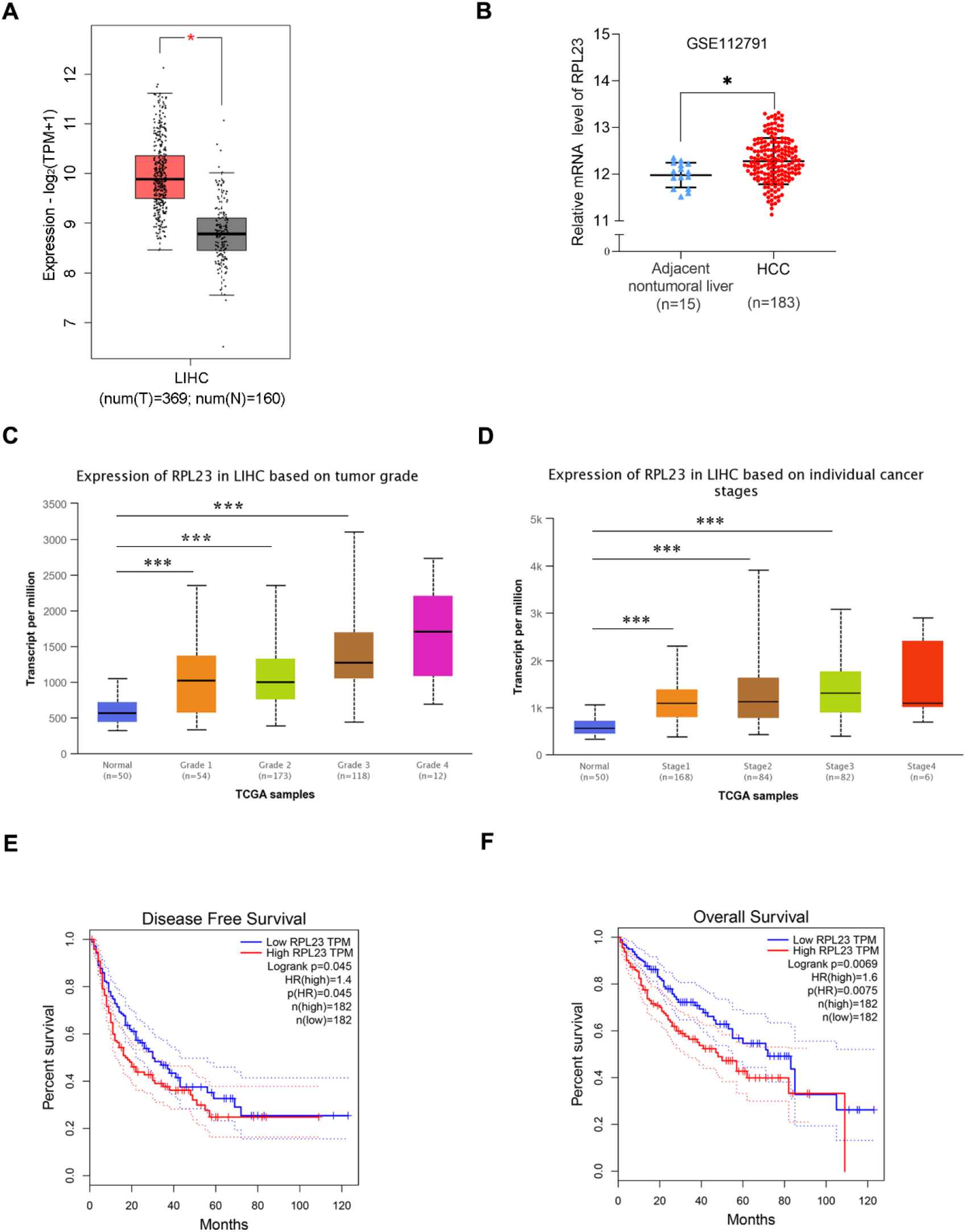
RPL23 overexpression and clinical pathological analysis in HCC tissues according to different public databases. (A) The mRNA level of RPL23 in the adjacent nontumor liver tissues (n=160) and primary liver tumor tissues (n=369) revealed by transcriptome sequencing of TCGA. The mRNA level of RPL23 from RNA-sequencing data on 183 primary human HCC tissues and 15 adjacent nontumor liver tissues of GEO database in the GSE112791 cohort. (C) RPL23 overexpression in HCC based on tumor grade in Ualcan data-mining platform (http://ualcan.path.uab.edu/index.html). (D) RPL23 expression in HCC based on tumor stage. (E) Correlation between RPL23 expression and disease-free survival in TCGA HCC cohort. (F) OS curve of HCC patients based on RPL23 expression in TCGA HCC cohort. *p<0.05, ***P < 0.001.

To further confirm the clinical significance of RPL23 expression in HCC, real-time PCR was performed to determine the mRNA level of RPL23 in human HCC tissues (T) and their adjacent nontumoral tissues (N) from 60 patients. The mRNA level of RPL23 was increased in human HCC tissues compared with their adjacent nontumoral tissues (Fig.2A), and the significant upregulation of RPL23 was observed in 87% of HCC tissues (Fig.2B). Consistently, the protein level of RPL23 was also increased in HCC tissue which assessed by Western blot and IHC (Fig.2C-D). Notably, the expression of RPL23 in EHMH was higher than that in MFH (Fig.2D). Moreover, the expression level of RPL23 was positively corelated to tumor vascular invasion (*p*=0.0070), lung metastasis (*p*=0.0469) and TNM stage (*p*=0.0346) in HCC (Table 1). In addition, the mRNA and protein level of RPL23 was dramatically increased in a panel of liver cancer cells compared to primary hepatocytes (PHH) (Fig.2E-F). Taken together, those data indicated that elevated RPL23 may involve in HCC metastasis and tumor progression.

**Figure 2.**
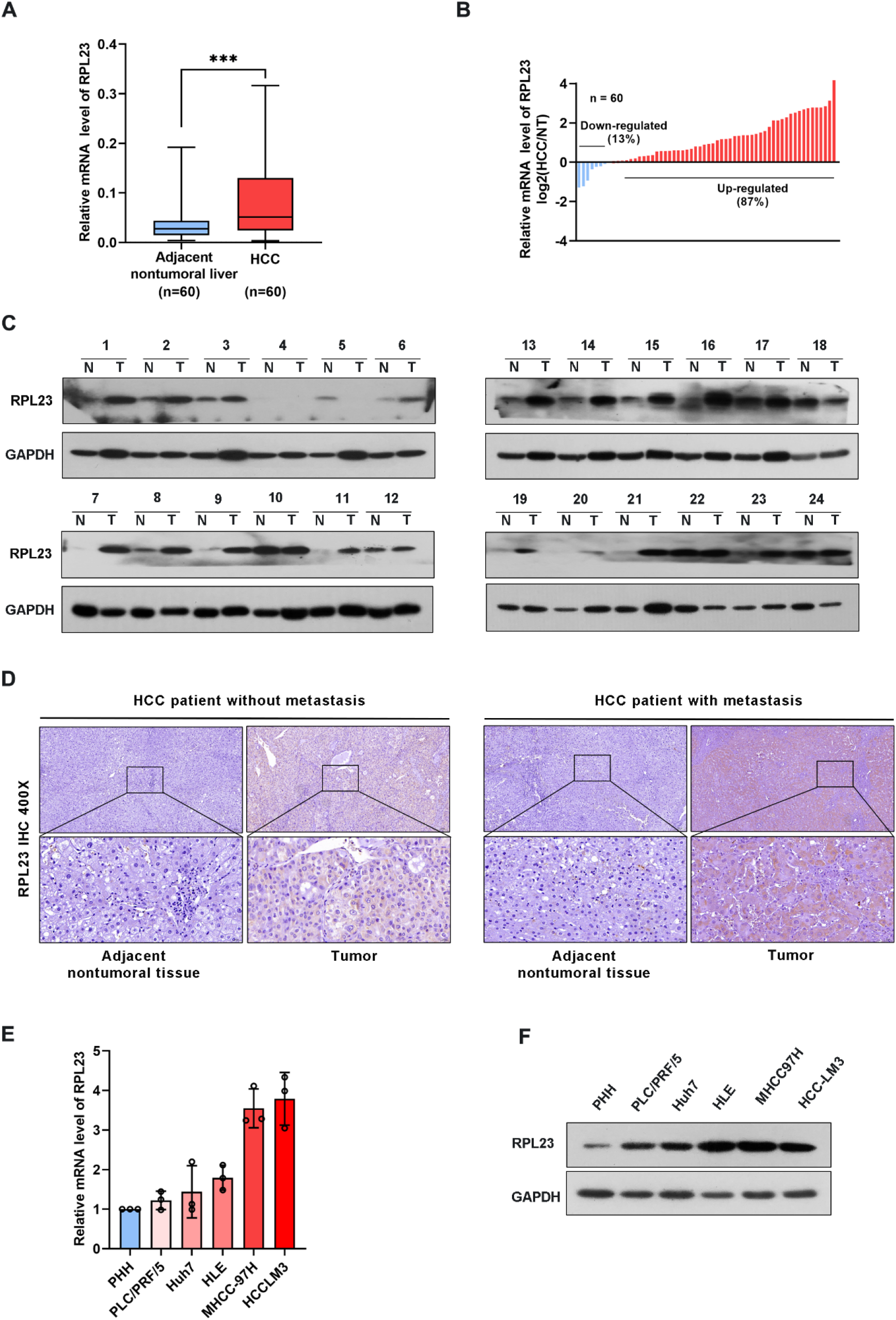
RPL23 levels increased in HCC tissues and overexpressed in different HCC cell lines. (A) The mRNA level of RPL23 in liver cancer was elevated in 60 paired samples of HCC tumor tissues with extrahepatic metastasis (EHMH) and metastasis-free HCC tissues (MFH)., as determined by qRT-PCR analyses. (B) RPL23 mRNA levels in 60 HCC and paired non-tumor tissues. (C) RPL23 protein level in 24 paired primary HCC tissues with extrahepatic metastasis (EHMH) and metastasis-free HCC tissues (MFH) was detected by western blotting. GAPDH was used as loading control. (D) Representative immunohistochemical images of RPL23 staining in the paired HCC specimens with or without metastasis. (magnification, 400×) (E, F) qRT-PCR and Western blotting analyses of RPL23 expression in primary human hepatocytes and different HCC cell lines. GAPDH was used as loading control. Representative data are from at least three independent experiments. Data are shown as mean ± SD. ***P < 0.001.

**Table 1.**
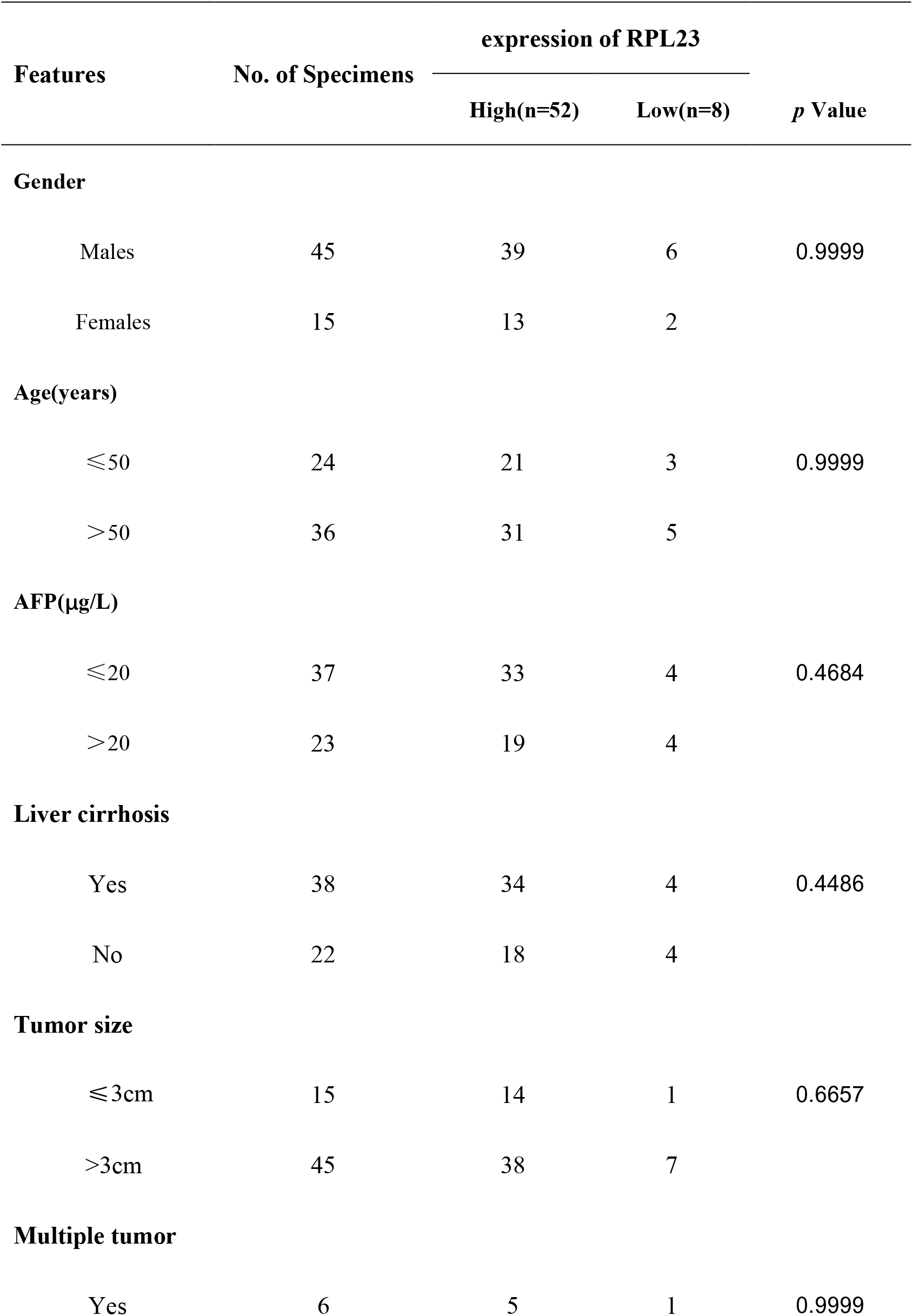
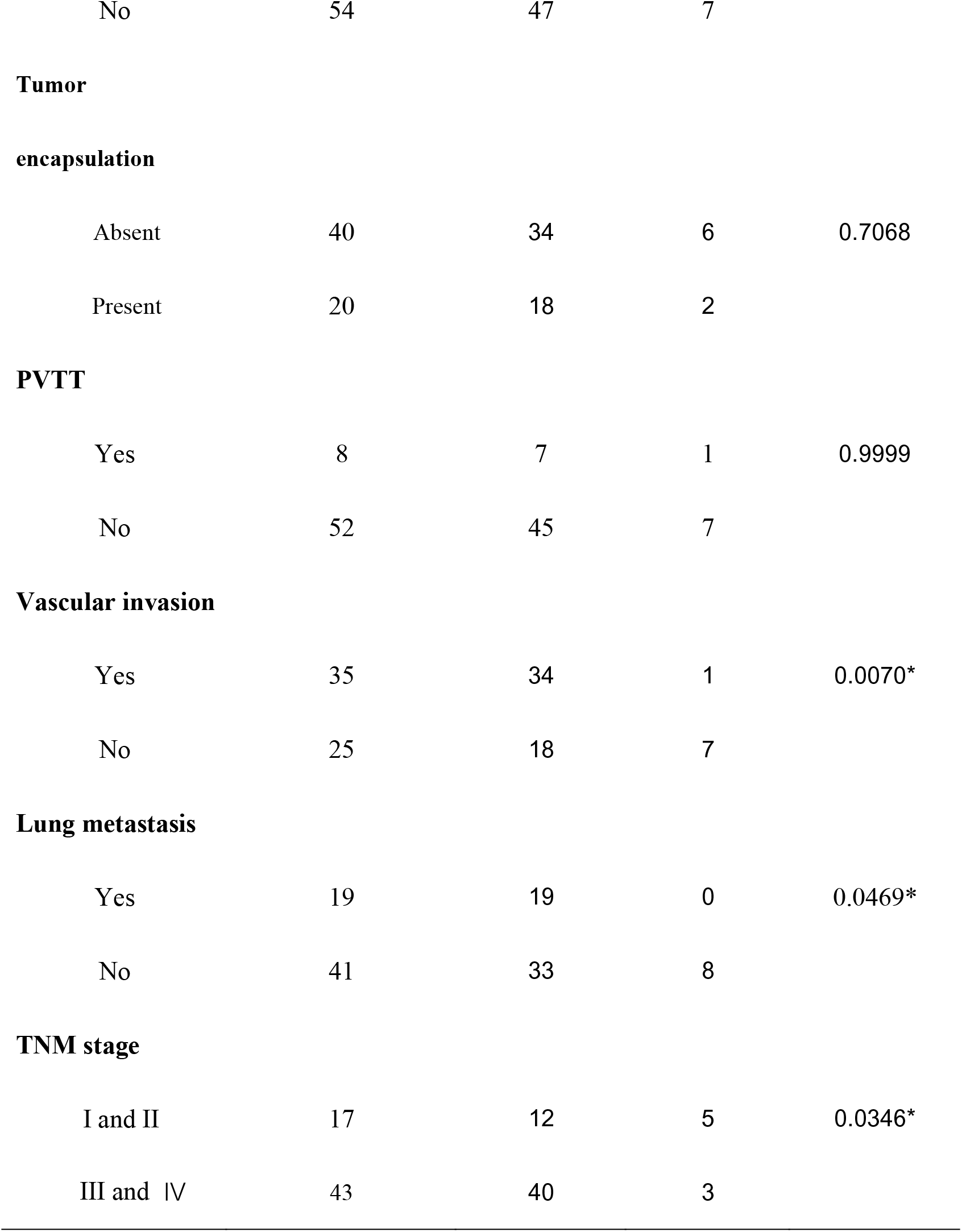
Correlation between RPL23 expression and clinicopathological characteristics in HCC patients

| Features | No. of Specimens | expression of RPL23 |  | *p* Value |
| --- | --- | --- | --- | --- |
|  |  | High(n=52) | Low(n=8) |  |
| Gender |  |  |  |  |
| Males | 45 | 39 | 6 | 0.9999 |
| Females | 15 | 13 | 2 |  |
| Age(years) |  |  |  |  |
| ≤50 | 24 | 21 | 3 | 0.9999 |
| >50 | 36 | 31 | 5 |  |
| AFP(μg/L) |  |  |  |  |
| ≤20 | 37 | 33 | 4 | 0.4684 |
| >20 | 23 | 19 | 4 |  |
| Liver cirrhosis |  |  |  |  |
| Yes | 38 | 34 | 4 | 0.4486 |
| No | 22 | 18 | 4 |  |
| Tumor size |  |  |  |  |
| ≤3cm | 15 | 14 | 1 | 0.6657 |
| >3cm | 45 | 38 | 7 |  |
| Multiple tumor |  |  |  |  |
| Yes | 6 | 5 | 1 | 0.9999 |
| No | 54 | 47 | 7 |  |
| **Tumor** |  |  |  |  |
| **encapsulation** |  |  |  |  |
| Absent | 40 | 34 | 6 | 0.7068 |
| Present | 20 | 18 | 2 |  |
| **PVTT** |  |  |  |  |
| Yes | 8 | 7 | 1 | 0.9999 |
| No | 52 | 45 | 7 |  |
| **Vascular invasion** |  |  |  |  |
| Yes | 35 | 34 | 1 | 0.0070* |
| No | 25 | 18 | 7 |  |
| **Lung metastasis** |  |  |  |  |
| Yes | 19 | 19 | 0 | 0.0469* |
| No | 41 | 33 | 8 |  |
| **TNM stage** |  |  |  |  |
| I and II | 17 | 12 | 5 | 0.0346* |
| III and IV | 43 | 40 | 3 |  |

### RPL23 silencing inhibited HCC cells proliferation, migration and invasion *in vitro*

To systemically evaluate the functions of RPL23 in HCC, we first examined the effect of RPL23 silencing on HCC cell growth and invasion. We chose HLE and MHCC97H cells, which express relatively high level of RPL23 compared with other HCC cell lines. The HLE and MHCC97H cells which stable expression of short hairpin RNA targeting RPL23 were generated and the efficiency was confirmed by Western blot (Fig.3A). Comparison with control cells, knockdown of RPL23 could decrease the proliferation rate in HCC cells (Fig. 3B). Moreover, cell migration was decreased by RPL23 depletion which witnessed by wound-healing assay (Fig.3C). Furthermore, transwell assay also confirmed that RPL23 depletion abolished the capacity to migrate and invade HLE and MHCC97H cells (Fig. 3D). Those data suggested that RPL23 may participate in HCC metastasis. It has reported that cytoskeleton remodeling mediated by actin filaments plays an important role in cell metastasis^24^ and therefore we examined the formation of actin filaments by using phalloidin staining. The data showed that RPL23 silencing led to the loose of actin filaments compared with the control cells (Fig. 3E), which benefit to cell migration and invasion. Overall, these data demonstrated that knockdown of RPL23 could inhibit HCC progression by repressing cell proliferation, migration and invasion.

**Figure 3.**
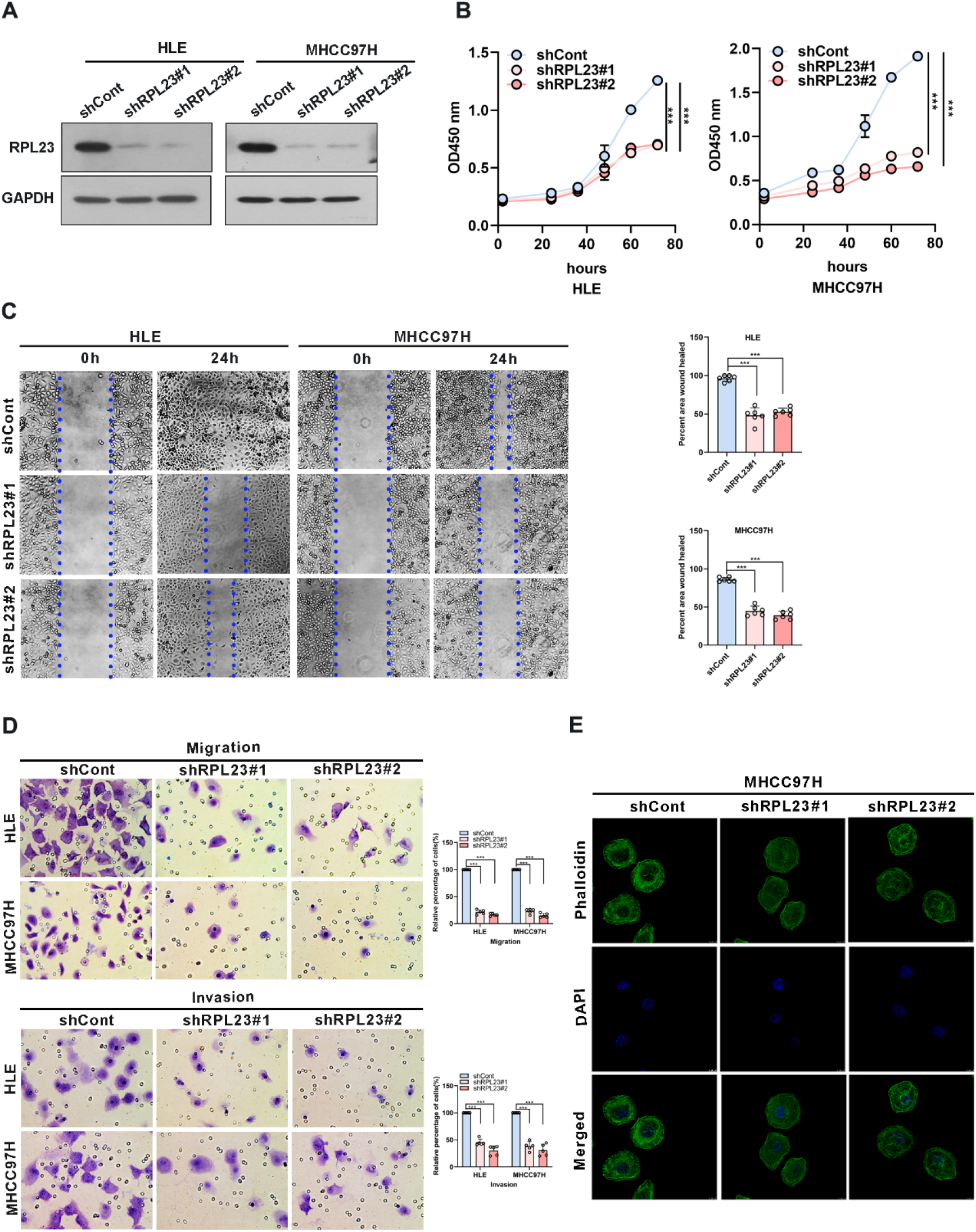
Knockdown of RPL23 significantly suppressed HCC cell proliferation, invasion and migration in vitro. (A) Western blotting was used to access RPL23 expression after transfected with negative control (shCont) or shRNA (shRPL23#1 and shRPL23#2) in HLE and MHCC97H cells. GAPDH was used as the internal quantitative control. (B) CCK-8 assay showed that RPL23 knockdown suppressed HCC proliferation capacity. (C) Wound-healing assays were performed to determine the migratory abilities of RPL23-knockdown HCC cells in HLE and MHCC97H cells. The cells were counted from 6 images. (D) Cell migration and invasion as measured by transwell assays were inhibited by knockdown RPL23 in in HLE and MHCC97H cells. The cells were counted from 5 images. (E) Phalloidin (green color) was applied for cytoskeleton staining, while DAPI (blue color) was used to mark the nuclei in RPL23-depleted MHCC97H cells. Magnification, 630×. Representative data are from at least three independent experiments. Data are shown as mean ± SD. ***P < 0.001.

### RPL23 overexpression promoted HCC cells proliferation, migration and invasion *in vitro*

To further confirm the biological role of RPL23 in HCC tumorigenesis, Huh7 cells which stable expression of RPL23 were constructed (Fig. 4A) and the effect of RPL23 overexpression on HCC progression were detected by series of experiments. Consistent with the findings in RPL23 depletion cells, cell proliferation rate was increased in RPL23 overexpression cells which determined by CCK-8 assay (Fig. 4B). Next, the wound-healing assay was conducted to determine the effect of RPL23 on cell migration. Aa expected, overexpression of RPL23 facilitate cell migration obviously (Fig. 4C). In addition, RPL23 overexpression cells were subjected to transwell assay and we found that ectopic expression of RPL23 promoted HCC cell migration and invasion (Fig. 4D). Collectively, these data indicated that overexpression of RPL23 could enhance the ability of HCC proliferation, migration and invasion.

**Figure 4.**
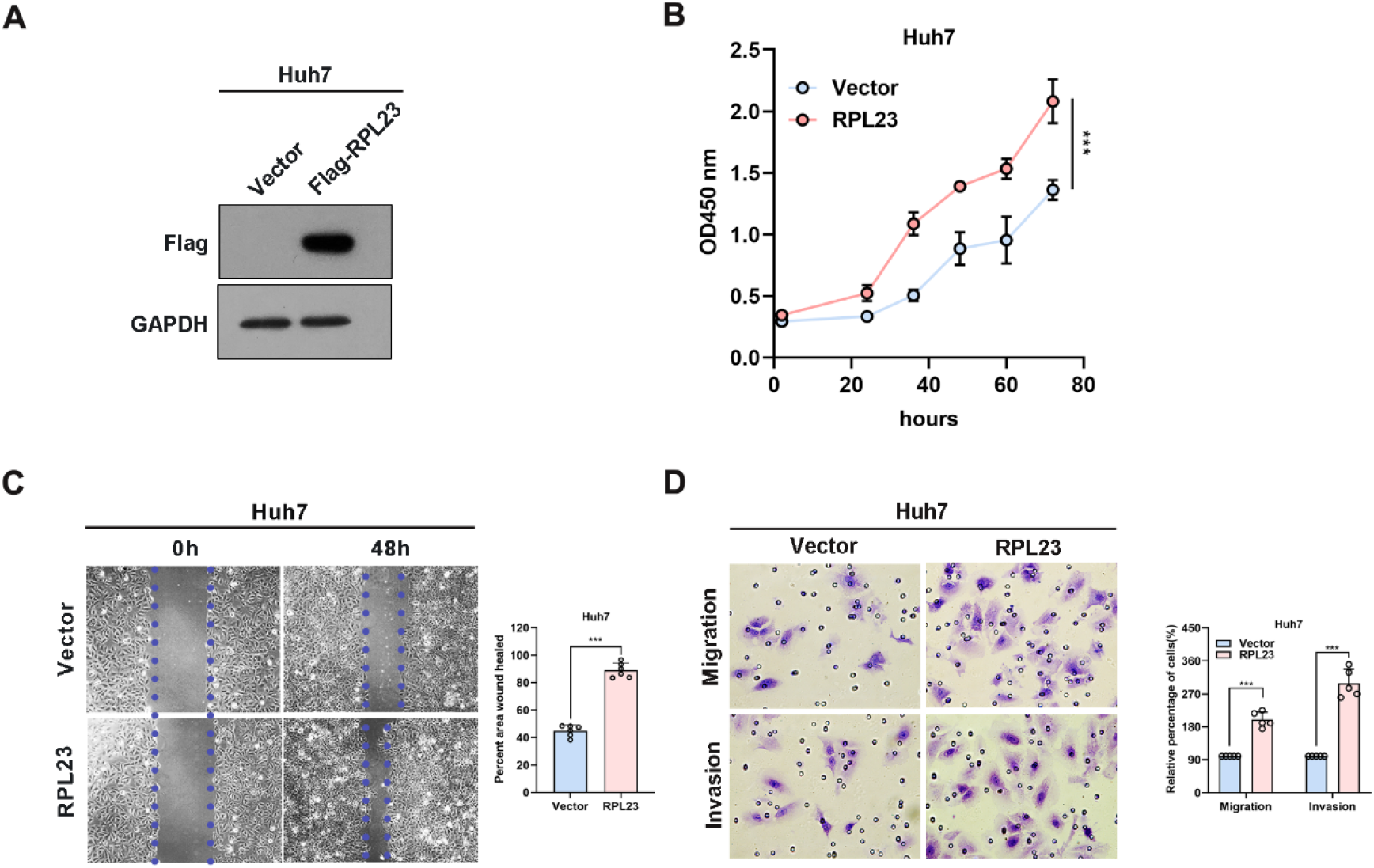
Overexpression of RPL23 promoted HCC cell proliferation, invasion and migration in vitro. (A) The overexpression efficiency of RPL23 was tested by western blotting analysis in Huh7 cells. GAPDH was used as a reference gene. (B) CCK-8 assay was applied to evaluate proliferation ability of Huh7 cells with expression of RPL23. (C) Migratory ability of overexpression RPL23 was detected by wound-healing assay in Huh7 cells. The cells were counted from 6 images. (D) Migratory and invasive ability were assessed by transwell assays in Huh7 cells with RPL23 overexpression. The cells were counted from 5 images. Representative data are from at least three independent experiments. Data are shown as mean ± SD. ***P < 0.001.

### RPL23 facilitates HCC metastasis via enhance MMP9 mRNA stability

Epithelial-mesenchymal transition (EMT) is a process whereby epithelial cells acquire mesenchymal features, resulting in decreased adhesion and enhanced migration or invasion. Considering that HCC cell migration and invasion could be enhanced by RPL23, we next analyzed the effect of RPL23 on EMT-associated markers in HCC cells. Based on real-time PCR data, we observed that RPL23 depletion could suppress MMP9 and MMP2 expression, while has no significant effect on other EMT-associated markers, such as N-cadherin, E-cadherin, Vimentin, Smad2 and Twist1 (Fig. 5A and 5B). Due to that MMP9 is the most significant downregulated genes by RPL23 silencing, the protein level of MMP9 were further analyzed by Western blot. The results demonstrated that RPL23 silencing lead to a remarkable decrease of MMP9 protein level (Fig. 5C). On the contrary, when RPL23 was overexpressed in Huh7 cells, expression of MMP9 was increased compared to the control cells (Supplementary Fig. 1A).

**Figure 5.**
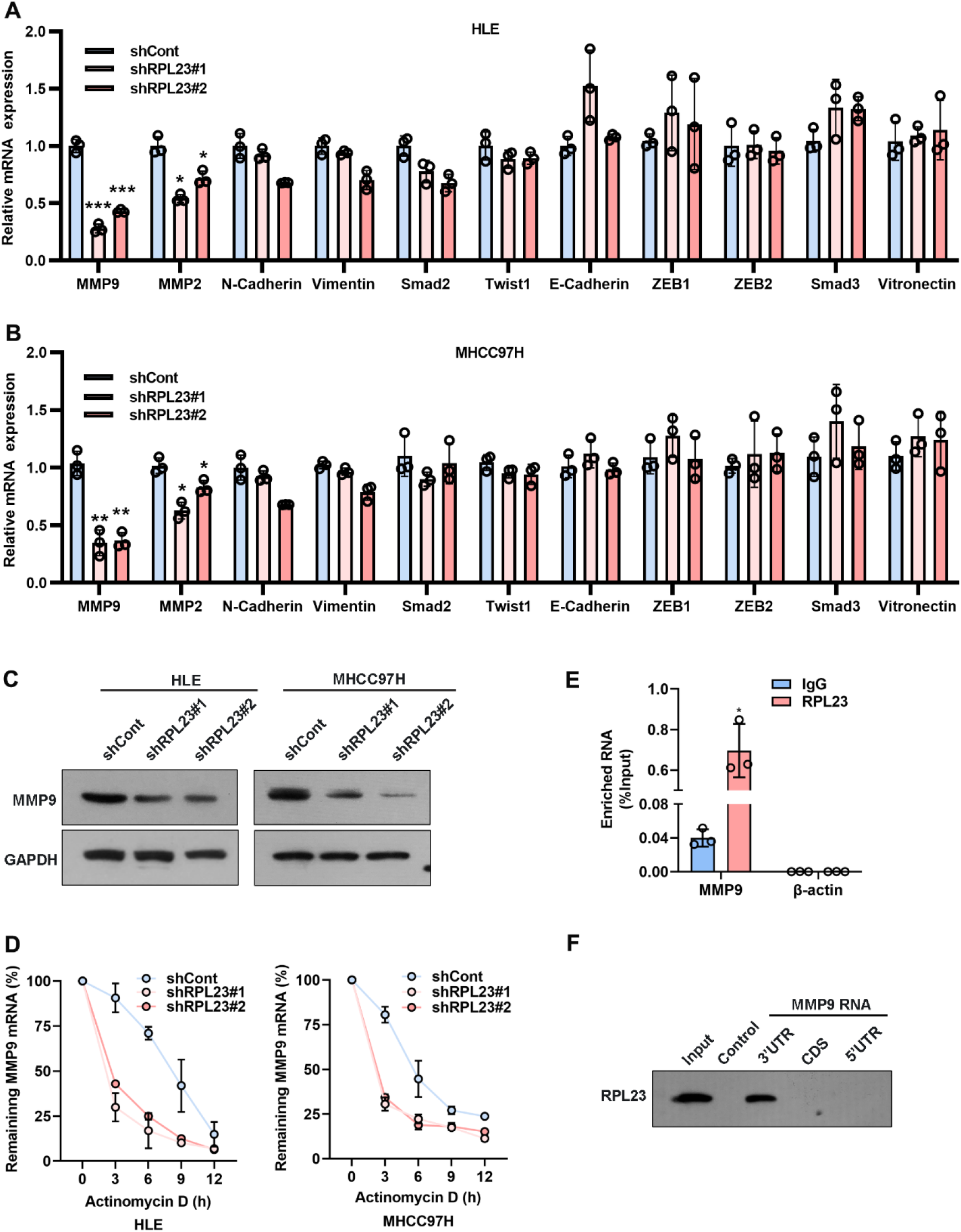
MMP9 is an essential downstream effector of RPL23. (A, B) EMT-related markers (MMP9, MMP2, N-cadherin, Vimentin, Smad2, Twist1 and E-cadherin) were measured on RPL23-depleted HCC cells by qRT-PCR. β-actin was used as an internal quantitative control. (***p<0.001) (C) RPL23 regulated MMP9 protein expression in HLE and MHCC97H cells measured by western blot assay. GAPDH was used as a loading control for western blotting. (D) The half-life of MMP-9 mRNA was reduced after RPL23 knockdown in HLE and MHCC97H cells followed by treatment with 5ug/mL actinomycin D at the indicated times. Error bars represent SEM. p-values (HLE): **p = 0.00116 (shCont vs shRPL23#1), **p = 0.00296 (shCont vs shRPL23#2). p-values (MHCC97H): **p = 0.00314 (shCont vs shRPL23#1), **p = 0.00477 (shCont vs shRPL23#2). (E) RIP assays showed that RPL23 directly bound to MMP9 mRNA. (F) RNA pull-down results showed that RPL23 was directly associated with the 3’UTR of MMP9 mRNA. Control indicates a control pulldown containing beads only. Representative data are from at least three independent experiments. Data are shown as mean ± SD. *p<0.05, **P < 0.01, ***P < 0.001.

Given that RPL23 is an RNA-binding protein which can regulate human cancer progress by influencing RNA stability ^25^, we speculated that RPL23 may regulate MMP9 mRNA level by modulating its stabilization. To determine whether RPL23 affects MMP9 mRNA stability in HCC cells, we treated RPL23 knockdown cells with actinomycin D to block transcription and measured decay of existing mRNAs by performing time-course real-time PCR. The results revealed that RPL23 depletion could shorten the half-life of MMP9 mRNA in RPL23 knockdown cells (t_1/2_=3 hours) compared to control cells (t_1/2_=5 hours) (Fig. 5D). Concertedly, RPL23 overexpression could delay the degradation of MMP9 mRNA in HCC cells (Supplementary Fig. 1B). To further reveal the direct association between RPL23 and MMP9, we performed a RNA immunoprecipitation (RIP) assay to assess whether RPL23 directly binds to MMP9 transcripts. The results demonstrated that MMP9 mRNA was significantly enriched in RPL23-IP sample compared with IgG-IP sample (Fig. 5E). Indeed, RNA pull-down results showed that RPL23 was directly associated with the 3’UTR of MMP9 mRNA (Fig. 5F). Taken together, our data provide a strong evidence that RPL23 promotes HCC metastasis by stabilization MMP9 mRNA via direct binding with the 3’UTR of MMP9.

To further study the functional role of MMP9 in RPL23-mediated HCC metastasis, we overexpressed MMP9 in RPL23 silencing cells and the effect on cell growth, migration and invasion were examined. Undoubtedly, the results showed that MMP9 overexpression phenocopied RPL23-induced cell proliferation (Fig.6A and 6B), and the inhibition of cell migration caused by RPL23 absence was rescued by MMP9 overexpression (Fig. 6C). Moreover, transwell assay confirmed that the effect of RPL23 depletion on cell migration and invasion were abolished by MMP9 (Fig. 6D). In short, we revealed that RPL23 could facilitate HCC metastasis in an MMP9 dependent manner.

**Figure 6.**
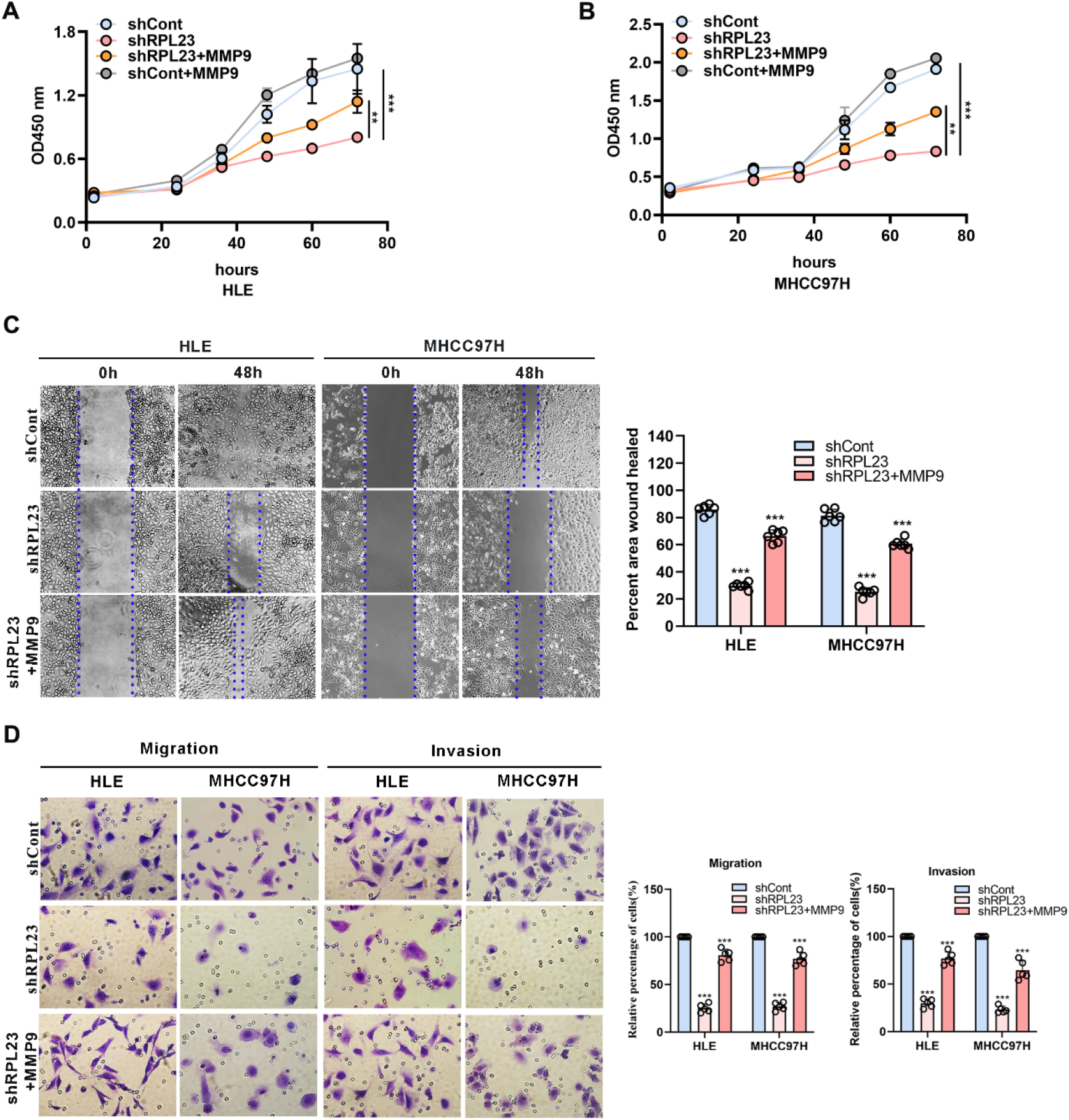
MMP9 overexpression rescues the RPL23 knockdown-induced malignant phenotypes. (A, B) Overexpression of MMP9 rescued the inhibition effect of decreased RPL23 on HCC cell proliferation. (C) overexpression of MMP-9 rescued the repression effect of knockdown RPL23 on HCC cell migration ability by wound-healing assay. The cells were counted from 6 images. (D) Upregulation of MMP-9 could significantly rescued the effects of decreased RPL23 in HLE and MHCC97H cells for both migration and invasion by transwell assays. The cells were counted from 5 images. Representative data are from at least three independent experiments. Data are shown as mean ± SD. **P < 0.01, ***P < 0.001.

**Figure 7.**
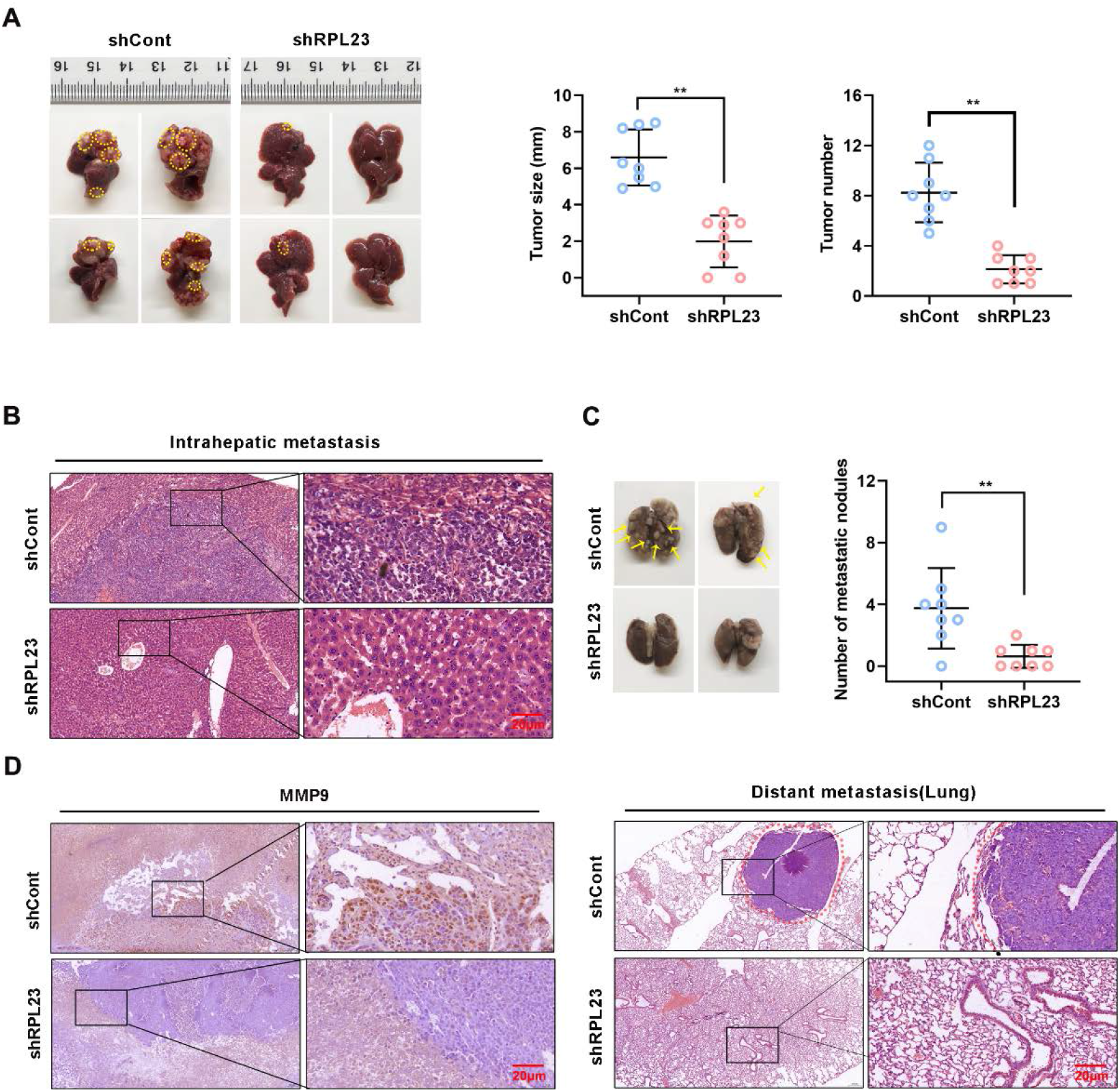
RPL23 knockdown suppressed HCC cell lung metastasis in vivo. The nude mice were orthotopically injected with MHCC97H cells stably depleted RPL23. (A) Representative images (left), volume and number (right) of xenograft liver tumor in nude mice. (B) The presence or absence of metastatic nodules in the liver was evaluated by Hematoxylin-Eosin staining. (C) Representative images of lung metastasis nodules in different groups (up), and evaluated by Hematoxylin-Eosin staining(down). (E) IHC assay showed MMP9 expression in metastatic xenograft model tissue in different groups. For A and C, the data were presented as mean ± SD, n = 8 in each group.

### RPL23 depletion inhibited HCC metastasis *in vivo*

To explore the inhibitory effect of RPL23 depletion on HCC metastasis *in vivo*, MHCC97H cells which stable expression of short hairpin RNA targeting RPL23 were injected into the left lobe of node mouse to generate the orthotopic nude mouse HCC model. Post 6-weeks injection, the mice were sacrificed and subjected to series detection. Consistent with *in vitro* observation, knockdown of RPL23 significantly decreased both the tumor growth rate and tumor size in liver (Fig. 6A and B). Moreover, the numbers of lung metastases in RPL23-depletion mice were significantly lower compared with those of control mice (Fig. 6C). Mechanistically, immunohistochemistry assay showed that MMP9 expression was apparently decreased in the liver of RPL23 depletion mice (Fig. 6D). In summary, those data strongly indicated that RPL23 depletion could downregulation the expression of MMP9 and therefore represses HCC metastasis.

## Discussion

Metastasis and postoperative recurrence are major contributors to the high mortality rate of HCC patients, which severely limit the improvement of clinical efficacy^26^. Therefore, actively exploring novel prognostic biomarkers and effective therapeutic targets of HCC recurrence and metastasis will be of great significance. It is well known that multiple levels of regulation are involved in the mechanism of occurrence and development of HCC metastasis, including transcriptional and post-transcriptional events. Of these, RBPs have received growing interest due to their critical roles in post-transcriptional control of mRNA by recognizing and binding to target sequences. Several previous studies showed that alterations in the expression and function of RBPs in HCC could amplify the effects of cancer driver genes, accelerate tumor progression, and promote tumor metastasis ^8^. Up to now, about 1542 human RBPs have been experimentally validated to be involved in diverse physiological and pathological processes including cancers ^5^. However, the amount of well-characterized RBPs involved in HCC is just the tip of the iceberg, and a great deal of HCC-related RBPs remain to be explored. In this study, we first identified that RPL23 promoted HCC tumorigenesis and metastasis via post-transcriptional regulation of MMP9 expression as an RBP in HCC.

Recent researches have uncovered that RPL23 is involved in various physiological and pathological processes, including cell proliferationc, cell apoptosis and cycle arrest. For example, in higher-risk myelodysplastic syndrome (MDS) patients, studies showed that overexpression of RPL23 was associated with the abnormal apoptotic resistance in CD34+ cells ^16, 27^. Watanabe et al. demonstrated that GRWD1 expression reversed the RPL23-mediated inhibition of anchorage-independent growth in HCT116 cells by negatively regulating RPL23 via the ubiquitin-proteasome system^28^. Meng and co-workers confirmed that RPL23-MDM2-p53 pathway could coordinate with the p19ARF-MDM2-p53 pathway against oncogenic RAS-induced tumorigenesis^29^. However, much of those literatures on RPL23 focus particularly on its effect in cell apoptosis, the underlying functions and mechanisms of RPL23 in HCC metastasis has been underestimated. In the current study, our data revealed that RPL23 was upregulated in HCC samples compared with adjacent normal tissues, and its high expression was associated with poor clinicopathological characteristics of HCC patients. Further analysis in experimental ex vivo and in vivo showed that knockdown of RPL23 inhibited the ability of proliferation, invasion and metastasis of HCC cells. Taken together, we proposed that RPL23, which was upregulated in HCC and significantly correlated with poor survival of HCC, could be a novel prognostic factor and potential therapeutic target for HCC.

Many alterations in tumor cells have been identified as contributing factors to the tumor metastasis. Among which, EMT is considered a key step promoting tumor invasion and metastasis for the following reasons: first, EMT can change the characteristics of tumor cells and therefore to lead to loss of junction molecules expression, reduction of the polarity and adhesion ability, and promotion of cell mobility, finally resulting in the enhancement of invasion ability of tumor cell; second, EMT can improve the ability of tumor invasion and metastasis by altering the microenvironment of tumor growth, invasion and metastasis and angiogenesis^30^; third, EMT can also increase the expression of several lytic enzymes which are responsible for the degradation of the extracellular matrix. As a result, tumor cells are more susceptible to shedding from primary sites^31^. Thus, to further explore the molecular mechanisms by which RPL23 promotes metastasis in HCC, we first turn our attention to the relationship between RPL23 and EMT associated genes. Our results found that the expression of MMP9 was markedly decreased after RPL23 silencing, and upregulation of MMP9 could markedly increase the weakened migration and invasion abilities of HCC cells induced by RPL23 knockdown, suggesting that MMP9 might be a potential downstream target of RPL23. Several literature reports have indicate that MMP9 plays a major role in migration of cancer cells^32^. Consequently, our data suggests a potential therapeutic approach to inhibit HCC metastasis by targeting the RPL23/MMP9 pathway.

A recent review reported by Pereira et al. comprehensively summarized the drivers mechanisms of RBPs in tumorigenesis and progression of tumors which included alternative splicing, polyadenylation, mRNA stability, subcellular localization and translation^5^. Zhao et al. reported that RPS3, a newly defined pro-tumorigenic RBP in HCC, contributed to the high level of the oncogene SITRT1 via stabilizing SIRT1 mRNA by binding to its 3’UTR in HCC cells ^33^.

Based on the observations mentioned above, we speculated that RPL23 might affect MMP9 expression by regulating its mRNA stability. Then this hypothesis was further confirmed by our experiments. We revealed that RPL23 up-regulated MMP9 expression by stabilizing MMP9 mRNA via binding to its 3’UTR. However, our findings about the interaction between RPL23 and MMP9 are still preliminary and deeper investigations are needed.

In summary, we report for the first time that RPL23 overexpression is a novel biomarker and prognostic factor for HCC. Mechanically, RPL23 promotes the expression of MMP9 by stabilizing MMP9 mRNA by binding to its 3’UTR, thus leading to a pro-metastasis effect in HCC. Our finding high light a possible new HCC treatment strategy that targets RPL23/MMP9 axis.

## Authors contributions

MLY, YJZ, HJD, FL and JC designed the study. MLY, YJZ, HJD, HZZ, STC, DPZ and XH performed the experiments and analyses. LM and YC provided the materials. MLY, YJZ and STC wrote the manuscript. FL and JC critically reviewed the manuscript. FL and JC supervised the study.

## Data availability statement

Additional supporting data are available from the corresponding authors on request. All requests for raw and analyzed data and materials will be reviewed by the corresponding authors to verify whether the request is subject to any intellectual property or confidentiality obligations.

## Supplementary

**Supplementary Figure 1.**
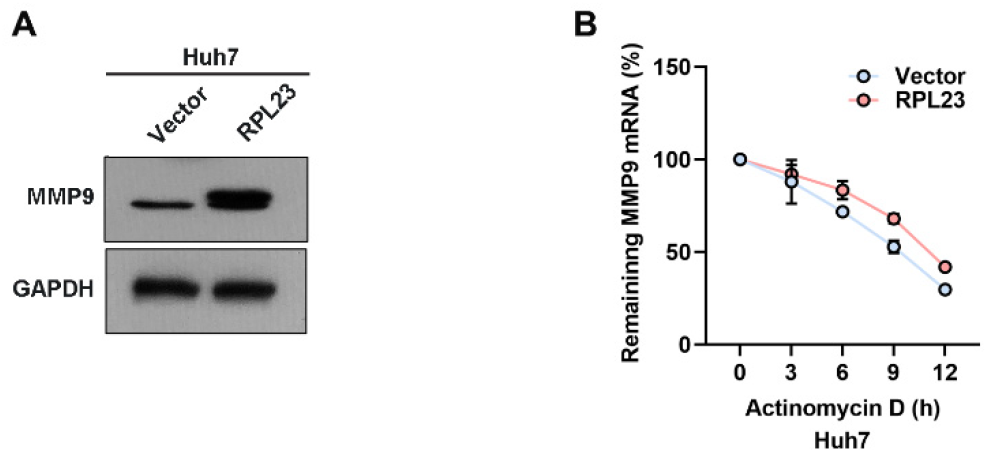
MMP9 is an essential downstream effector of RPL23. (A) The protein level of MMP9 increased after overexpression of RPL23 in Huh7 cells. The half-life of MMP-9 mRNA was increased after RPL23 expression in Huh7 cells followed by treatment with 5ug/mL actinomycin D at the indicated times. *p = 0.0187 (Vector vs RPL23). Representative data are from at least three independent experiments. Data are shown as mean ± SD.*p<0.05.

**Supplementary table 1:**
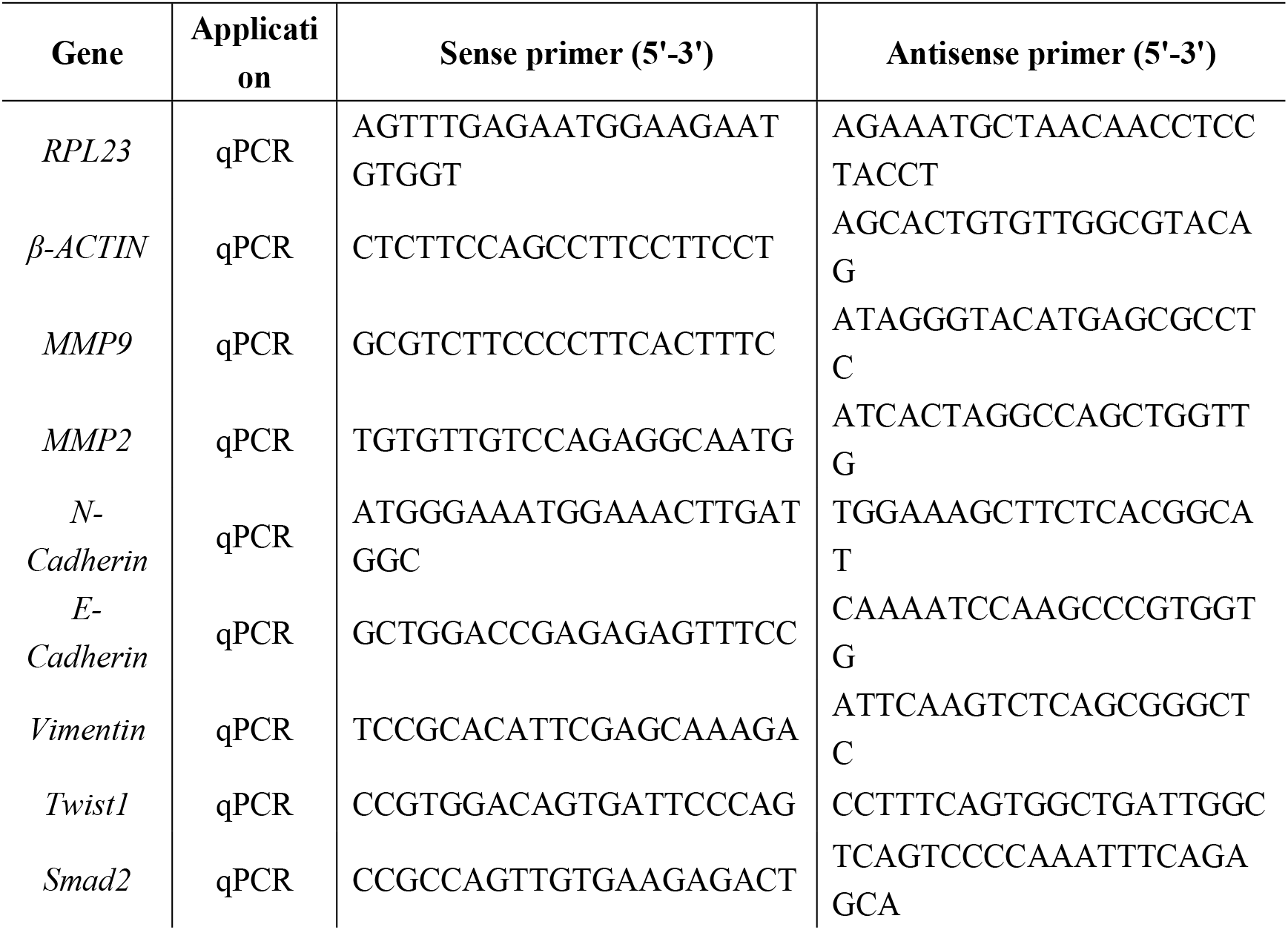
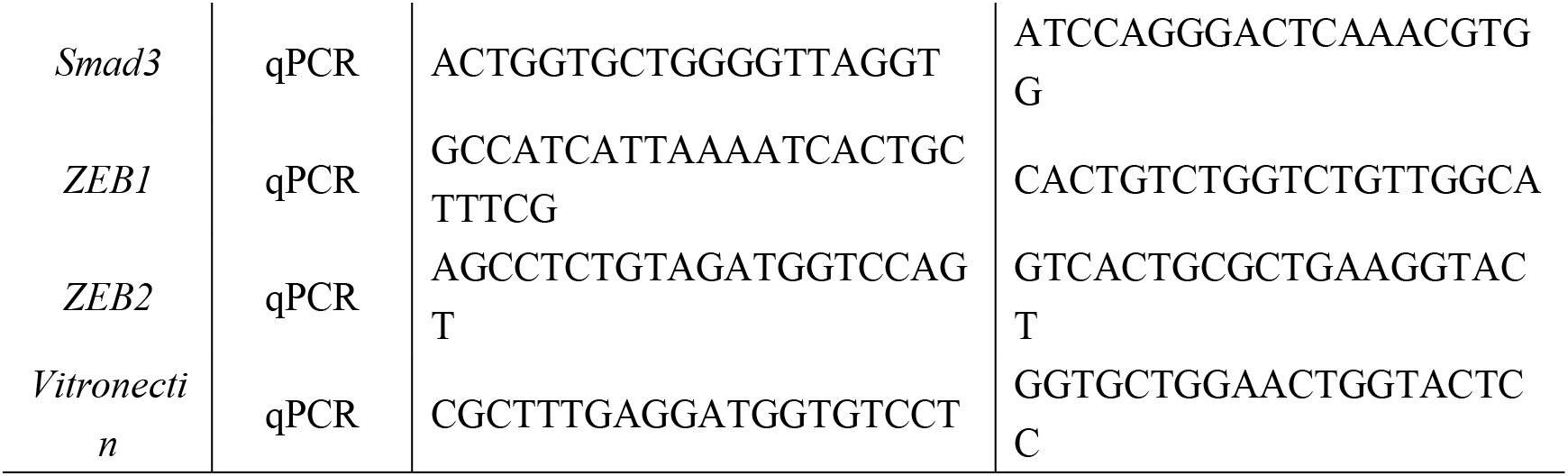
DNA oligonucleotides used for qPCR

